# Loss of *Fgfr1* and *Fgfr2* in Scleraxis-lineage cells leads to enlarged bone eminences and attachment cell death

**DOI:** 10.1101/2021.09.20.461087

**Authors:** Kendra K. Wernlé, Michael A. Sonnenfelt, Connor C. Leek, Elahe Ganji, Zachary Tata, Anna Lia Sullivan, Elijah Paparella, Claudia Offutt, Jordan Shuff, David M. Ornitz, Megan L. Killian

**Author notes:** Corresponding author: Megan L. Killian, PhD. Author contributions: Conceptualization: Lead- DMO and MLK; Supporting- KKW and MAS Data curation: Lead- KKW, MAS, and MLK; Supporting- CCL, EG, ALS, ZT Formal analysis: Lead- KKW, MAS, CCL, MLK; Supporting- EG, ZT, EP, CO, JS Funding acquisition: Lead- MLK; Supporting- DMO Investigation: Lead- KKW, MAS, CCL, MLK; Supporting- EG, ZT, ALS, EP, CO, JS Methodology: Lead- KKW, MAS, CCL, MLK; Supporting- EG, ZT, ALS, DMO Project administration: Lead- KKW, MLS, MLK Resources: Lead- MLK Software: Lead- KKW, MAS, CCL, MLK Supervision: Lead- MLK Validation: Lead- KKW, MAS, CCL, MLK Visualization: Lead- KKW, MAS, CCL, MLK; Supporting- DMO, EG, EP, ALS, CO, ZT Writing- original draft: Lead- KKW, MAS, MLK; Supporting- CCL, EG, ZT, DMO Writing- review and editing: Lead- KKW, MAS, MLK; Supporting- CCL, EG, ZT, ALS, EP, CO, JS, DMO.

## Abstract

Tendons and ligaments are structural tissues that attach to bone and are essential for joint mobility and stability in vertebrates. Tendon and ligament attachments (i.e., entheses) are often found at bony protrusions (i.e., eminences), and the shape and size of these protrusions depends on both mechanical forces and cellular cues during growth and development. The formation of tendon eminences also contributes to mechanical leverage for skeletal muscle. Fibroblast growth factor receptor (FGFR) signaling plays a critical role in bone development, and *Fgfr1* and *Fgfr2* are highly expressed in the perichondrium and periosteum of bone where tendon and ligament attachments can be found. However, the role of FGFR signaling in attachment development and maintenance in the limb remains unknown. In this study, we used transgenic mouse models for combinatorial knockout of *Fgfr1* and/or *Fgfr2* in tendon/ligament and attachment progenitors using ScxCre and measured eminence size and bone shape in the appendicular skeleton. Conditional deletion of both, but not individual, *Fgfr1* and *Fgfr2* in *Scx* progenitors led to enlarged eminences in the postnatal appendicular skeleton and smaller secondary ossification centers in long bones. In addition, *Fgfr1 Fgfr2* double conditional knockout mice had more variation in the size of collagen fibrils in tendon, narrowed synovial joint spacing, and increased cell death at sites of ligament attachments, as well as decreased plasticity of mature bone compared to age-matched wildtype littermates. These findings identify a role for FGFR signaling in regulating growth and maintenance of tendon/ligament attachments and the size and shape of bony eminences.

## Introduction

Tendons and ligaments transmit mechanical forces to bone and are essential for joint mobility and stability in vertebrates (1–3). During development, the attachments of most tendons and ligaments of the appendicular skeleton mature into graded fibrocartilage interfaces between tendon/ligament and bone (2,4–6). These fibrocartilage attachments are often found at bone eminences, which are superstructures such as tuberosities and tubercles, that increase leverage between the line of action of applied mechanical force (e.g., skeletal muscle) and the center of a joint (7–9). In mice, eminence formation initiates around embryonic day (E)12.5 from a recruited progenitor pool that co-expresses *Scleraxis (Scx*) and *Sox9* and surrounds the cartilage anlage of developing bone (7,10). Eminences form “miniature” growth plates at tendon/ligament attachments and follow similar developmental patterns to the primary growth plate during longitudinal bone growth (1,8,9,11–14). In the absence of muscle loading, such as paralysis, chondrocyte proliferation within the mini-growth plate of the developing eminence is arrested (8). In addition, the absence of muscle loading also leads to malformed synovial joints in embryonic limbs (15–22). However, it remains unclear if and how growth and shape of tendon and ligament attachments are regulated by similar molecular cues to the growth plate.

Fibroblast growth factors (FGFs) play a critical role in bone development and maturation (23). FGF ligands bind with high affinity to transmembrane tyrosine kinase receptors, FGFRs, and these ligands and receptors directly interact with heparan sulfate proteoglycans (23), which are richly deposited during cartilage development as well as at sites of tendon and ligament attachments (24). Recent work has demonstrated cell-specific mechanisms of both *Fgfr1* and *Fgfr2* signaling in long bone development (25,26). Expression of *Fgfr1* is localized to the hypertrophic chondrocytes of the developing growth plate, and both *Fgfr1* and *Fgfr2* are highly expressed in the perichondrium and periosteum (23). Loss of *Fgfr1* and *Fgfr2* in osteoprogenitor cells results in increased expression of *Fgfr3* and *Fgf9* and the non-autonomous regulation of growth plate chondrocytes, leading to suppressed chondrocyte proliferation and shorter bones (25). In mice lacking *Fgfr1* expression in mature osteoblasts and osteocytes, the osteocytes die resulting in secondary bone overgrowth (26). Surprisingly, although global deletion of *Fgf9* leads to shortened bones as well, developing eminences of the humerus (e.g., deltoid tuberosity) becomes noticeably enlarged (27,28). This differential regulation of growth between long bone growth plates and eminence “mini” growth plates led us to further explore the role that FGFR signaling plays in the size and shape of tuberosities. Recently, work by Roberts et al. showed that deletion of *Fgfr2* in neural crest cells leads to smaller eminences in the craniofacial complex (29). Yet the link between FGFR signaling and postnatal eminence maturation in the appendicular skeleton remains unclear.

In this study, we show that eminence growth and maintenance in the mammalian limb is regulated in part by FGFR signaling. Loss of *Fgfr1* and *Fgfr2* in tendon/ligament progenitors led to unrestricted growth of eminences in the appendicular skeleton, increased cell death at ligament attachments, reduced synovial joint spacing, and reduced plasticity of long bones. These results identify a critical role of FGFR signaling in regulating growth and maintenance of tendon/ligament attachments at eminences for controlling the shape and structure of bones.

## Materials and Methods

### Animals

All animal use was approved by the University of Delaware Institutional Animal Care and Use Committee and Washington University Animal Care and Use Committee. Wild-type (WT) and *Fgfr* conditional knockout (KO) mice were bred in-house on a mixed C57BL/6J; 129X1 background in a specific pathogen-free facility and handled in accordance with standard use protocols and animal welfare regulations. A total of 79 mice were used for this study. Sample size of experimental and control groups varied based on breeding outcomes: *Fgfr1*^flx/flx^*;Fgfr2*^flx/flx^ mice were crossed with ScxCre;*Fgfr1*^flx/WT^; *Fgfr2*^flx/WT^ mice to generate *ScxCre;Fgfr1*^flx/flx^;*Fgfr2*^flx/flx^ mutants (DKO; n = 24; n = 3 at postnatal day 1 [P1]; n = 7 at 3-4 weeks of age; n = 5 at 8 weeks of age; and n = 9 at 6-8 mo of age), as well as single mutants of *ScxCre;Fgfr1*^flx/flx^(R1cKO, n=4 at 6-8 mo of age) and *ScxCre;Fgfr2*^flx/flx^ (R2cKO, n=2 at 6-8 mo of age). Cre-negative littermates were used as WT controls (n = 28; n = 6 at P1; n = 8 at 3-4 weeks of age; n = 6 at 8 weeks of age; and n = 8 at 6-8 mo of age). Homozygous *ROSA^tdTomato^*(Ai14) reporter mice (Madisen, 2010, PMID: 20023653) (Jackson Laboratory, Bar Harbor, Maine) were crossed with ScxCre mice to generate ScxCre; *ROSA^tdTomato^* reporter mice (n = 20).

### Lineage tracing

Knees of ScxCre; *ROSA^tdTomato^* reporter mice were dissected at the time of euthanasia (three time points: postnatal day 1-2; 2 weeks of age; or 2-3 mo of age) and fixed in 4% paraformaldehyde for lineage tracing experiments. Tissues were decalcified in 14% EDTA for 2 weeks, embedded in SCEM (Section-Labs, Japan) or OCT (Fisher Scientific), and sectioned at 10μm. Sections were stained with DAPI (Vector Labs, CA, USA) and imaged on an inverted epifluorescent microscope (Zeiss AxioObserver Z1, Thornwood, NY).

### Micro-computed tomography

Left hindlimbs and forelimbs of young adult mice at 8 weeks of age (n = 6 WT; n = 5 DKO) and adult mice at 6-8 mo of age (n = 5 WT; n = 6 DKO; n = 4 R1cKO; n = 2 R2cKO) were fixed in 4% paraformaldehyde for 24-48 hr with care taken to not disrupt the natural joint flexion angle. After unilateral limb dissection, carcasses were frozen at −20°C (non-defrost freezer) for long-term storage prior to mechanical testing. Limbs were scanned using microcomputed tomography (microCT; Scanco Micro35, Switzerland; 20μm voxel size, 55kV, 72μA, 300ms integration time). Tibial length and bone mineral density of the cortex, as well as humeral length, epiphyseal volume, and deltoid tuberosity volume (WT and DKO only), were measured for 8-week-old and 6-8 mo-old mice. DICOM stacks were reconstructed in three-dimensions using OsiriX MD (Pixmeo, Switzerland) for visualization of frontal and sagittal planes. Reconstructed microCT images of the knee were three-dimensionally rendered using OsiriX MD to measure joint angle and tibial slope. Lateral and medial meniscal thickness was measured using sphere-fitting algorithms (30) in CTAN (Bruker, Belgium). The unmineralized volume of the knee joint was measured using frontal-plane views of the knee between the tibial plateau and femoral condyles.

For bone shape analyses, long bone scans (from 8 week and 6-8-mo-old mice, WT and DKO only) were reconstructed and analyzed using Dragonfly software (Object Research Systems, Montreal, Quebec). Bones were aligned along their respective neutral axes. Once aligned, a “Box Form” was created around each long bone of interest. Using the “Extract Structural Grid” function, the bone of interest was realigned and extracted. From the realigned version of each long bone of interest, lengths were measured between the most proximal point of the proximal epiphysis and the most distal point of the distal epiphysis. Volumetric data was collected using a “clip box” generated in Dragonfly to isolate the deltoid tuberosity (DT), lateral supracondylar ridge (LSR), and entire humerus for each sample. A threshold was defined to include only bone voxels for the DT and LSR. The “Split at Otsu” function was used to extract bone voxels for the entire humerus. The “Bone Analysis” function in Dragonfly was then used to fill in the bone ROI and measure the total bone ridge volume.

### Histology

After microCT scans were complete, limbs were decalcified in formic acid or 14% EDTA (Immunocal, StatLab, TX, USA) for 72 hr or 2 weeks, respectively. Tissues were processed for paraffin sectioning, sectioned at 7μm, stained with Toluidine Blue (StatLab, Texas, United States), and imaged using an upright bright field microscope (Zeiss Axio Imager 2, Thornwood, NY). Supraspinatus attachment width was approximated using Toluidine Blue-stained histological sections using Image J (31). Histological sections from knees of 3-4 week old mice (WT and DKO groups; n=3/genotype) were used for terminal deoxynucleotidyl transferase dUTP nick end labeling (TUNEL) to visualize cell apoptosis/death (Roche, Basel, Switzerland). Sections were counterstained with DAPI and visualized using epifluorescence (Nikon Eclipse Ni E800). The total number of nuclei and TUNEL+ cells were counted in cropped regions of cruciate ligament-bone attachments in each knee; both caudal and cranial cruciate attachments on the femur and tibia, when visible, were measured using the particle analyzer tool using Fiji (32). At least two ligament attachments were measured for each biological replicate. The percentage of TUNEL+ cells was calculated for each attachment and percentages of TUNEL+ cells for each attachment were averaged within each knee.

### Transmission electron microscopy (TEM)

The Achilles tendons of adult WT and DKO mice (6-8 mo of age; n = 3 per genotype) were dissected and fixed for 2 weeks in a 2% glutaraldehyde, 2% paraformaldehyde solution buffered to pH 7.4 with 0.1M sodium cacodylate. Tendons were cut in the transverse plane from the proximal tendon and imaged to visualize collagen fiber diameters and tendon ultrastructure (Zeiss Libra 120 Transmission Electron Microscope; 120kV, 0.34nm point to point resolution, and Gatan Ultrascan 1000 CCD camera, Pleasanton, CA). Fiber area, fiber diameter, number of fibers, and fiber density were measured using automated segmentation of at least three (3) technical replicates per sample in Dragonfly (Object Research Systems, Montreal, Quebec). Only two WT tendons were included in the analysis (with at least 3 technical replicates for TEM sections) due to poor infiltration of fixative.

### Mechanical testing

Frozen carcasses from WT and DKO mice (6-8 mo of age; previously used for microCT and histology) were thawed for mechanical testing. Femurs were carefully dissected and stored at 4°C in phosphate-buffered saline (PBS) and tested within 24 hr post-dissection at room temperature. Femoral length was measured using a digital caliper. Three-point bending was performed using a custom-made fixture with 7mm span length and rounded top press-head supports (2 mm diameter) were used to minimize shear during compression. All tests were performed at room temperature in air. Bones were positioned horizontally with dorsal side facing up in the fixture. A 0.2 N preload was applied, and femora were tested quasi-statically under displacement control with a crosshead speed of 0.1 mm/sec to failure. All mechanical testing was performed on an electromechanical uniaxial tester (Instron 5943, Norwood, MA) with a 100N load cell and 10 Hz recording range. From load-displacement data, the following data were calculated using a custom MATLAB (MathWorks, Natick, MA) code: ultimate load, displacement at maximal force, post-yield displacement (displacement at fracture - displacement at maximal force), fracture displacement, stiffness, and work to fracture (area under curve of the load-displacement curve to failure). The contralateral femurs of WT and DKO mice, previously scanned using microCT, were evaluated as follows to determine long bone geometry: DICOM images were binarized and realigned about the neutral axis to obtain transverse image stacks using ImageJ (31). Area moment of inertia was then approximated about the minor axis (I_min_) using BoneJ (33).

### Statistical Analysis

All statistical comparisons, including descriptive statistics, and graphical representation of data were performed using Prism 8.4.3 or later versions (Graphpad Software LLC, San Diego, CA). Three-way ANOVAs (genotype, age, medial/lateral menisci) with repeated measures by region were used to compare meniscal thickness and joint volume within and between each group. For direct comparisons between WT and DKO mice, two-sample t-tests were used to compare biomechanical properties of bone. Two-way ANOVAs were used to compare epiphyseal volumes between WT and DKO mice at 8 weeks and 6 mo of age and a one-way ANOVA was used to compare flexion angles between genotypes at 6-8 mo of age. Two-way ANOVAs were used to compare tibial and humeral length between WT and DKO mice at 8 weeks and 6–8-mo time points. Due to low sample size for R2cKO (n = 2), statistical comparisons were used to compare WT and DKO groups only and not statistically compared between single FGFR mutants. TUNEL-positive cell ratios were compared using an unpaired t-test of within-sample averages.

## Results

### Scx-lineage targets tendon/ligament and attachments as well as the Groove of Ranvier

Lineage tracing with experiments showed consistent localization of ScxCre; *ROSA^tdTomato^*(Ai14) cells (*Scx*Cre tdTomato+ lineage) in tendons, ligaments, and entheses (Figure 1). Additionally, tdTomato+ cells were found within the secondary ossification center of bone (at ~3-4 weeks of age) and the mineralized zone of the growth plate, similar to the *Osx-Cre* lineage (34,35). TdTomato+ cells from the ScxCre; *ROSA^tdTomato^* lineage were prevalent in the perichondrium surrounding the proximal and distal sites of the secondary ossification centers, which differs from previous reports using lineage tracing of mesenchymal progenitors (e.g., *Osx-* Cre; Figure 1) (36). Also, unlike *Osx*-Cre lineage tracing, *Scx*Cre tdTomato+ cells were localized to tendon, ligament, articular cartilage, and other fibrous/fibrocartilaginous connective tissues (e.g., meniscus).

**Figure 1.**
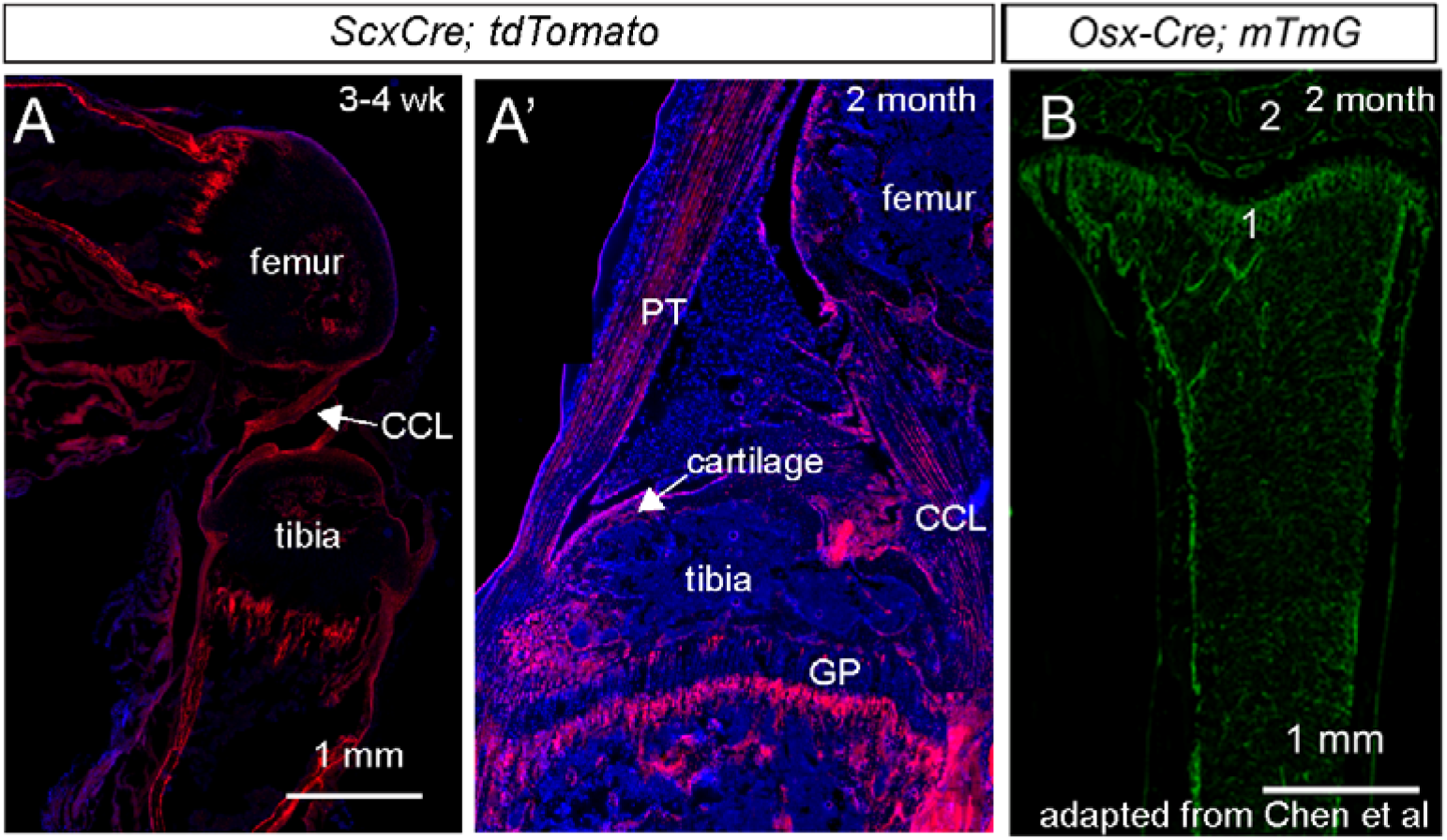
Lineage tracing of ScxCre. (A) Lineage tracing (ScxCre; *ROSA^tdTomato^* reporter) of ScxCre at 2weeks of age and (A’) 2mo of age. (B) Previously reported lineage tracing with Osx-Cre (Osx-Cre; *ROSA^mTmG^*, GFP reporter) at 2 mo of age (34). 1 = trabecular bone of the growth plate and 2 = secondary ossification center. CCL = cranial cruciate ligament; GP = growth plate; PT = patellar tendon.

### ScxCre-DKO mice had larger tuberosities, yet shorter bones, compared to WT littermates

There were no gross phenotypic differences at the time of weaning between WT and DKO mice; however, DKO mice weighed less (15.6 ± 2.34g) compared to WT mice (18.6 ± 1.94g; p<0.05) at 8 weeks of age. Additionally, DKO mice consistently had shorter bones compared to WT mice at 8 weeks and 6 mo of age (Figure 2A). In spite of DKO mice having shorter humeri, tuberosities on the humeri (the deltoid tuberosity, DT; and lateral supracondylar ridge, LSR) were significantly larger in DKO mice compared to WT mice at 6 mo of age (Figure 2B). This enlarged tuberosity phenotype was also observed at sites of tendon attachments in the tibia (Figure 3A).

**Figure 2.**
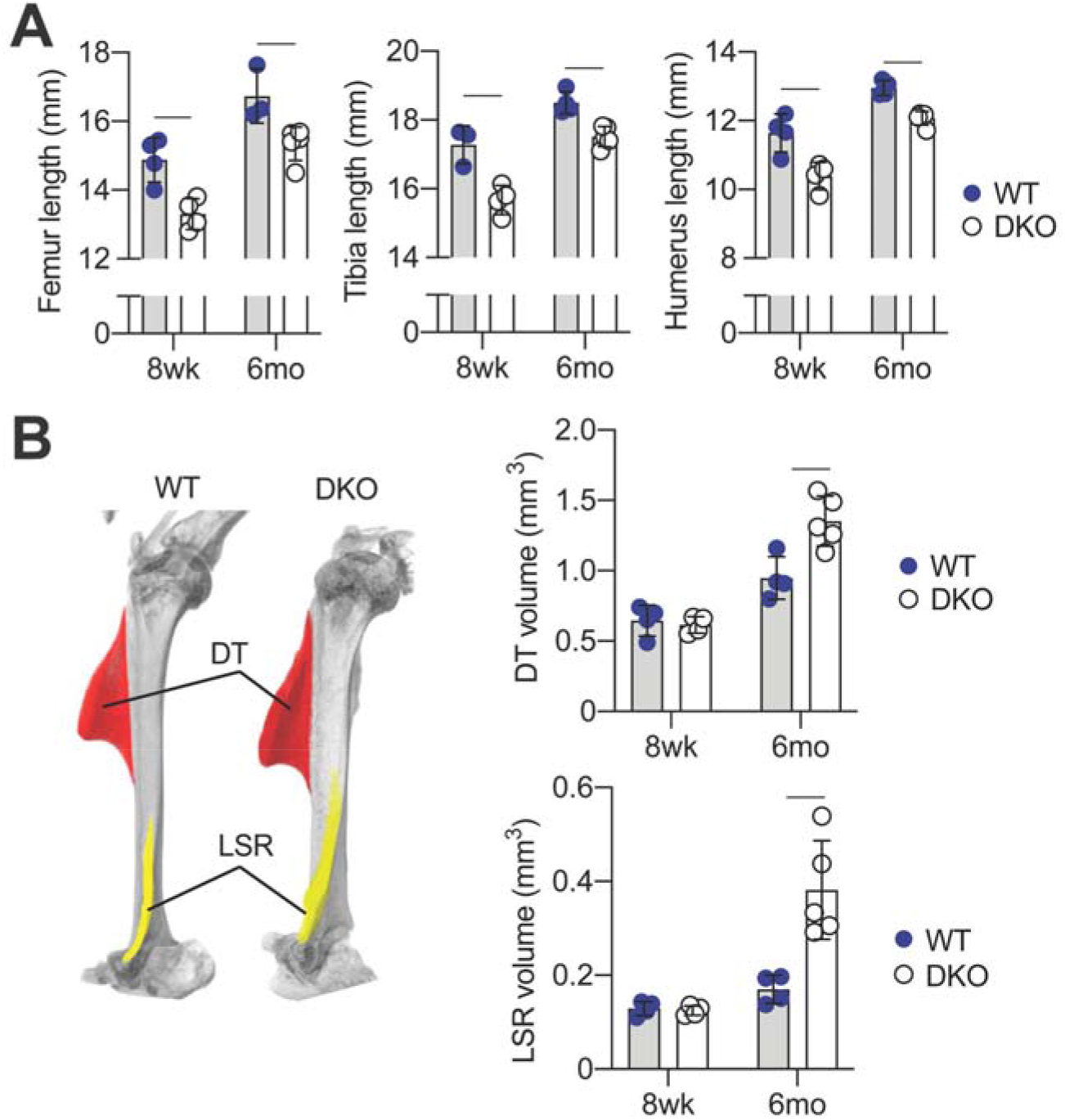
DKO mice had shorter bones with larger tuberosities. (A) Long bone lengths (femur, tibia, and humerus) in DKO mice were significantly shorter at 8 weeks and 6mo compared to age-matched WT mice. (B) Superstructure volumes, visualized in representative microCT reconstructions of 6mo old WT and DKO humeri, were significantly larger in DKO mice compared to age-matched WT mice. Pseudo-colored structures show the representative regions used to measure volumes of the deltoid tuberosity (DT) in red, and the lateral supracondylar ridge (LSR) in yellow.

**Figure 3.**
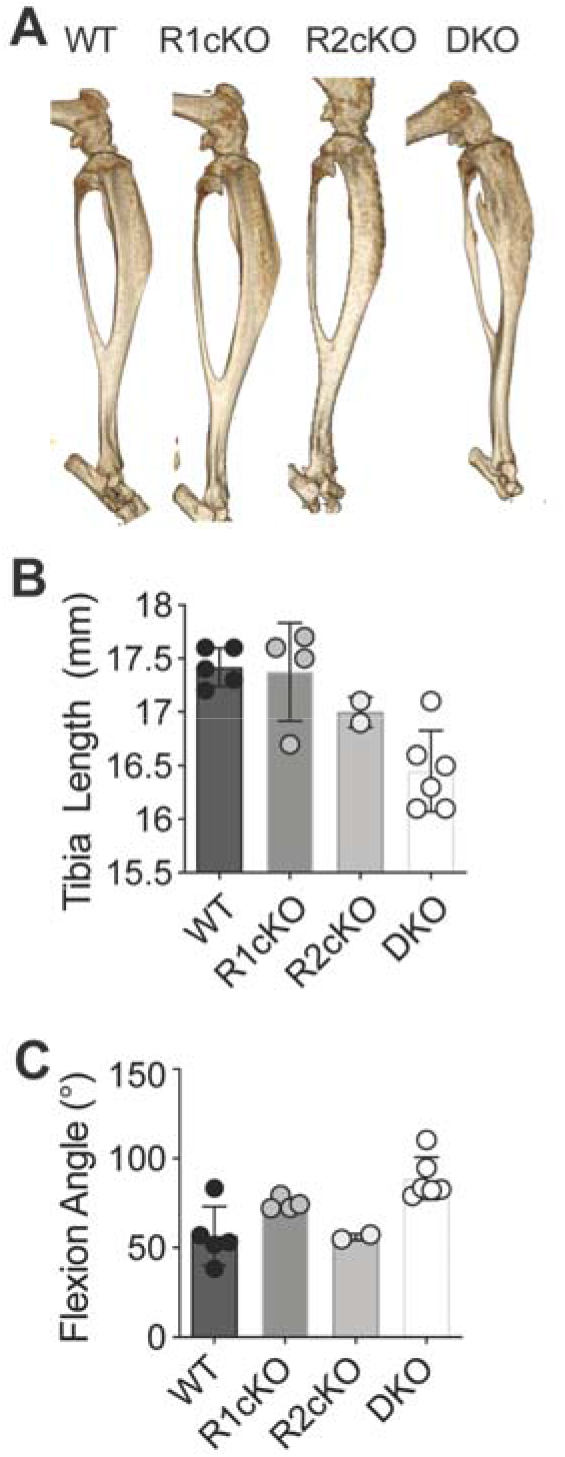
DKO mice had smaller tibias and increased knee flexion angles. A) MicroCT reconstructions of tibia from WT, *Fgfr1* cKO, *Fgfr2* cKO, and DKO mice at 6-8 mo of age. B) Tibia length was dependent on the number of alleles of *Fgfr1* and *Fgfr2* that were deleted. C) Flexion angle was greater in DKO mice compared to other genotypes.

### DKO mice had narrowed knee joint spacing and increased cell death at ligament entheses

Based on the distinct phenotype related to increased tuberosity size in adult DKO mice, we also investigated the shape and morphometry of articular joints, specifically the knee. DKO mice had greater knee flexion and reduced tibial slope compared to WT littermates at 6-8 mo (Figure 4A and B). Single receptor mutants (*Fgfr1* or *Fgfr2* cKO) did not have any remarkable bone phenotypes when compared to DKO mice, with comparable tibia lengths and flexion angles to WT mice at 6-8 mo of age (Figure 3B and C). Tibial cortical bone mineral density (BMD) was significantly lower in DKO mice (1,090 ± 66mg HA/cm^3^ ) compared to WT mice (1,167 ± 50mg HA/cm^3^) at 6-8 mo of age.

**Figure 4.**
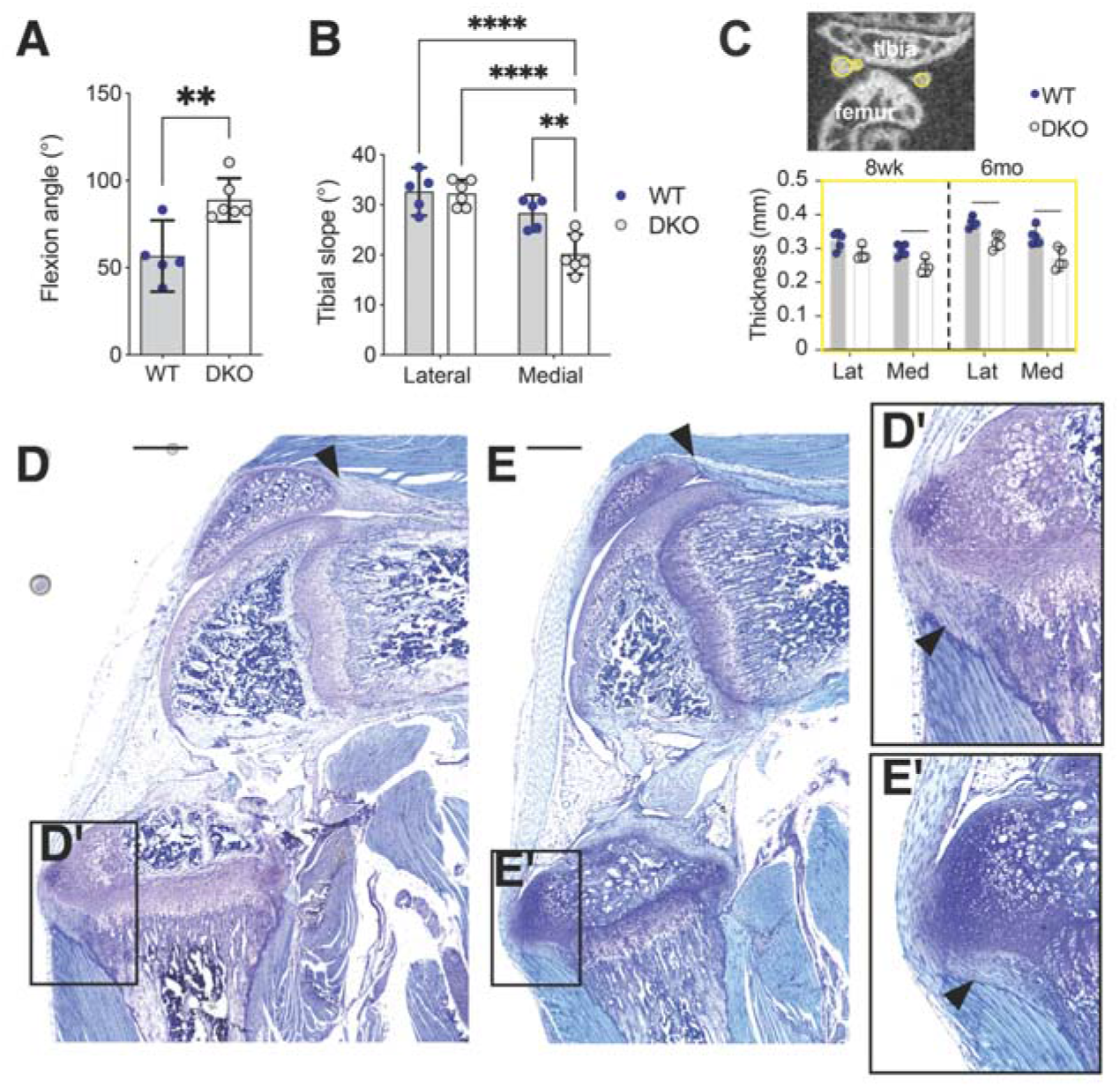
Decreased knee joint spacing and smaller Groove of Ranvier in DKO mice. (A) DKO mice had more knee flexion and (B) decreased medial tibial slope compared to WT mice at 6 mo of age. (C) Mineralized meniscal thickness was measured using sphere-fitting, shown in yellow, and the maximum diameter was used to estimate knee joint spacing thickness. At 8 weeks of age, DKO mice had thinner medial knee joint spacing, and by 6 mo of age, both lateral and medial joint spacing were thinner in DKO mice compared to WT mice. Knee morphology at 3 weeks of age for (D) WT and (E) DKO showed marked differences in the morphology of the quadriceps tendon (black arrows) and (D’, E’) the Groove of Ranvier, which were smaller in DKO mice. Scale bar = 100μm.

In mice, the meniscus is mineralized as early as 8 weeks of age; therefore, we used microCT to estimate joint space by measuring the thickness of the mineralized meniscus at 8 weeks and 6 mo of age (Figure 4C). DKO mice had significantly thinner knee spacing in the medial compartment at both 8 weeks and 6-8 mo of age compared to WT mice, and this thinning was also present in the lateral compartment at 6 mo of age in DKO mice (Figure 4C). Histologically, the patella and proximal patellar tendon appeared smaller in DKO mice compared to WT mice at 3 weeks of age (Figure 4D,E) as did the groove of Ranvier (Figure 4D’,E’).

Based on these morphological findings, we further investigated cell viability at the attachment site of the intraarticular ligaments of the knee. Attachment sites of cruciate ligaments in 3 week old DKO knees had significantly increased cell apoptosis, indicated by increased TUNEL+ staining, compared to attachment sites in age-matched WT knees (Figure 5). Apoptotic cells were primarily localized to the mineralizing region of the interface and were not found in the fibrous ligament tissue (Figure 5B, E, F). No other regions in histological sections of the mouse knee had a remarkable presentation of apoptotic cells.

**Figure 5.**
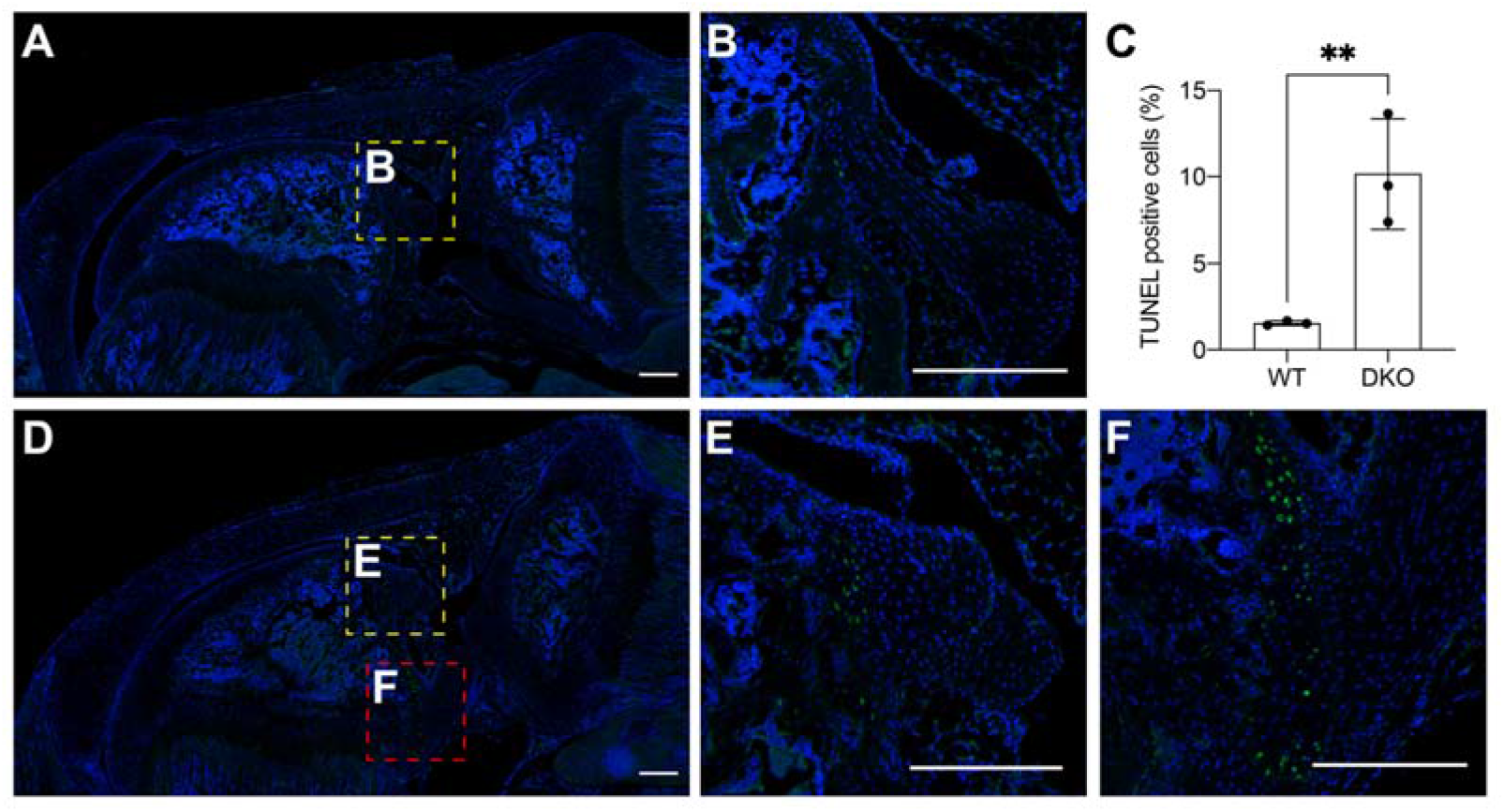
Increased cell death in DKO ligament entheses at 3 weeks of age. Cruciate ligament attachments of (A,B) WT knees had relatively low cell death, marked by TUNEL+ staining, compared to similar attachments in (D-F) DKO knees. (C) DKO ligament entheses had significantly higher rates of TUNEL+ cells compared to age-matched WT entheses. Scale bar = 250μm.

### Increased collagen fibril size in DKO compared to WT tendons

The ultrastructural properties of tendon collagen fibrils were analyzed in the Achilles tendon at 6-8 mo of age in WT and DKO mice (Figure 6A,B). DKO mice had a higher frequency of small fibrils in Achilles tendon compared to WT mice (~40-50nm diameter) and also had more larger collagen fibrils compared to WT (~260-300nm; Figure 6C). The relative area covered by collagen fibrils was higher in DKO tendons compared to WT tendons (Figure 6D), but the average fibril area was not significantly different between groups (Figure 6F).

**Figure 6.**
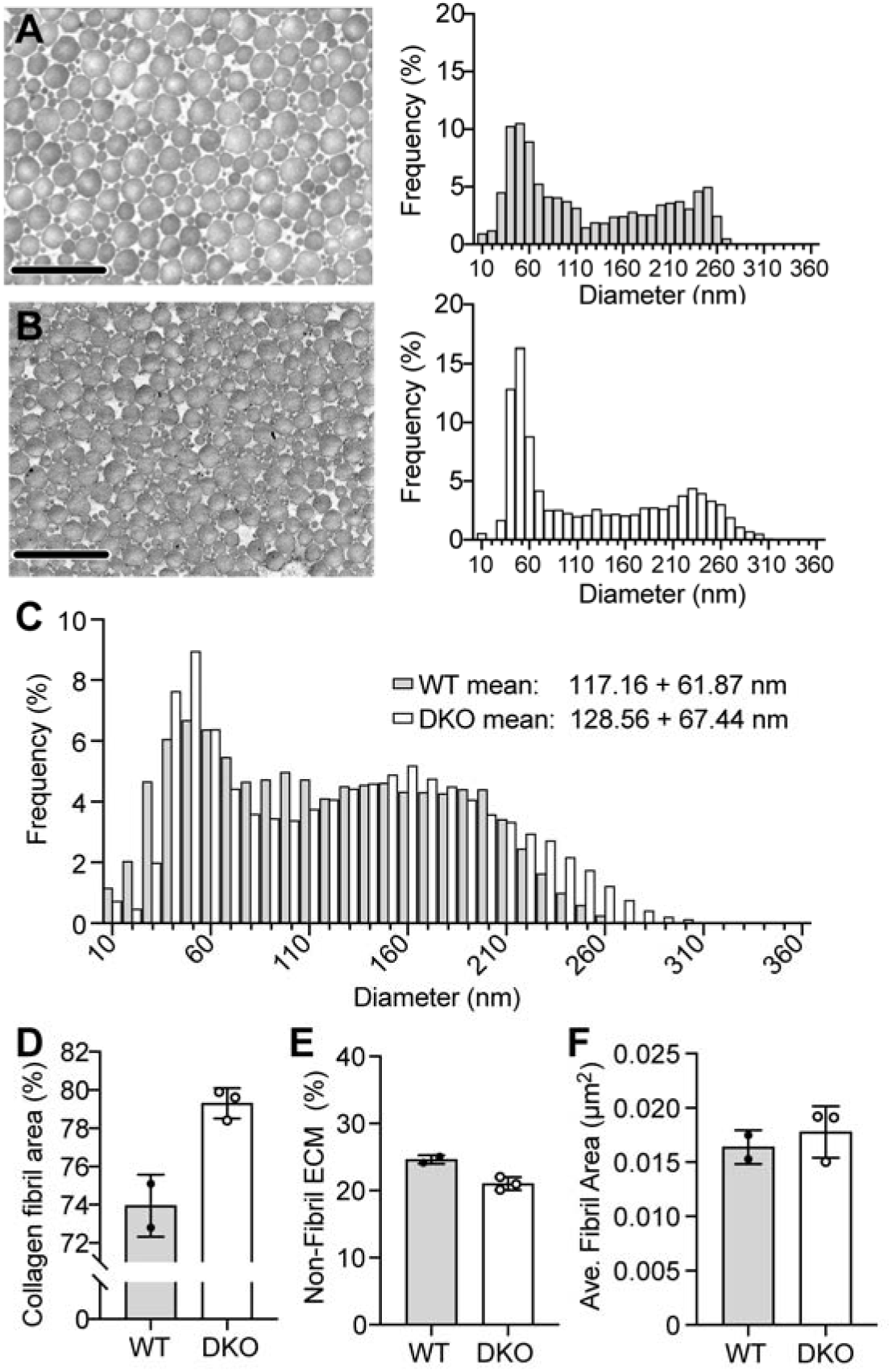
Collagen fibril ultrastructure of the Achilles tendons of WT and DKO mice. Transmission electron micrographs and representative histograms of frequency distribution of fibril diameter (10 nm increments) of (A) WT and (B) DKO tendons (scale bar = 1μm). (C) Overlaid histogram of representative histograms for WT and DKO mice with average fibril diameters and standard deviations. (D) Collagen fibril area (%) was calculated for field of views and (E) non-fibril extracellular matrix area was measured as the entire area of the field of view excluding collagen fibrils. (F) Average fibril area (μm^2^) was measured using a watershed function in Dragonfly.

### DKO long bones were more brittle compared to WT bones

Biomechanical properties of DKO and control bones were assessed by three point bending. The post-yield behavior of DKO femurs were impaired compared to WT at 6-8 mo of age (Figure 7A). Although the femur length was significantly shorter in DKO mice (Fig 7B), the ultimate load was not significantly different between groups (Figure 7C). Both post-yield displacement (Figure 7D) and work to fracture (Figure 7E) were significantly reduced in DKO femurs compared to WT femurs. Surprisingly, we did not find significant differences between genotypes in cross-sectional geometry of the femurs (moment of inertia; WT = 0.15 ± 0.04mm^4^ vs. DKO = 0.19 ± 0.04 mm^4^).

**Figure 7.**
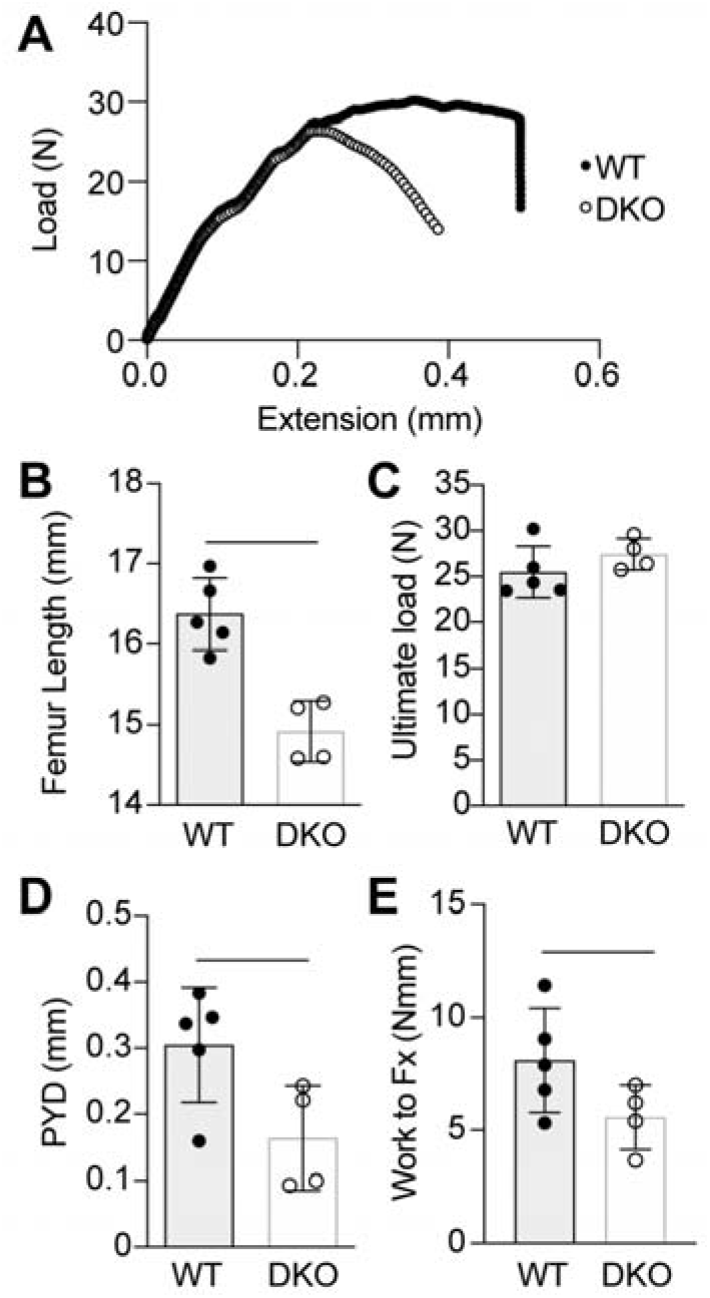
Whole bone biomechanical properties. (A) Representative load-extension curves for WT (black) and DKO (white) femurs at 6-8 mo of age. (B) Femur lengths (mm) were significantly shorter in DKO compared to WT. (C) Ultimate load (N) was not significantly different between groups, but (D) post-yield behavior (post-yield displacement, PYD); and (E) work to fracture (Work to Fx) were significantly reduced in DKO femurs compared to WT femurs.

## Discussion

Each bone in the adult mammalian skeleton has a unique shape and structure that accommodates its functional requirements for movement and stability. The formation of superstructures along the surface gives bone their topographical shape and provide distinct attachment sites for tendons, yet the mechanism by which these superstructures grow has only recently been explored (8). In this study, we showed that FGF signaling plays a role in regulating the shape and structure of these superstructures in the mature skeleton. We discovered that ablation of FGF signaling in developing tendon and attachments results in aberrant bone shape, such as enlarged mineralized superstructures, and also alters the ultrastructure of tendon collagen fibrils. These changes in the structural properties may influence functional outcomes, such as bone strength, and could also alter joint movement. From these studies, it remains unclear if the tendon ultrastructure was a primary outcome of altered FGF signaling or if these changes were caused by a secondary effect of altered tendon loading induced by irregular joint shape and more mechanistic studies of the developing tendon and enthesis are needed. Nonetheless, we found that adult mice lacking *Fgfr1* and *Fgfr2* in tendon and attachments had increased knee flexion and reduced knee joint space. Additionally, deletion of *Fgfr1* and *Fgfr2* in *Scx*-lineage cells led to impaired expansion of secondary ossification centers in long bones, resulting in shorter bones and flattened epiphyses. McKenzie et al. recently described increased osteocyte death in mice lacking *Fgfr1* and *Fgfr2* either in osteoblasts or osteocytes that led to increased cortical remodeling and bone accrual (26). We also found increased cell death in the subchondral bone of ligament attachments in the knees of young ScxCre-DKO mice, which may have directly or indirectly caused increased bone apposition at the enthesis. These cells may be derived from hedgehog-responsive enthesis progenitors (e.g., Gli1+) or epiphyseal chondrocyte (e.g., Col2+) cells (13). Further investigation into the mechanism by which enthesis progenitors are maintained and how FGF signaling regulates cell survival and bone accrual at the enthesis are needed.

Deficiencies in functional behavior in ScxCre-DKO bones, especially decreased postyield displacement, suggest collagen-level (nanoscale) changes in bone plasticity. In connective tissue (i.e., tendon), ultrastructural analysis showed that some DKO collagen fibrils were larger and their size distribution differed compared to normal collagen fibrils in WT tendons. The size and structure of collagen fibrils in tendon is controlled by small leucine-rich proteoglycans (e.g., decorin) containing glycosaminoglycan chains of either chondroitin sulfate or dermatan sulfate. Ablation of decorin and its complimentary protein, biglycan, results in larger fibril diameter in tendons, similar to what we observed in ScxCre-DKO mice (37). Heparan sulfate proteoglycans (HSPG), which preferentially bind FGF ligands and receptors, may also regulate type I collagen fibrillogenesis (38,39), and additional work is needed to explore how HSPG regulates collagen fibril formation in both tendon and bone. Additionally, the differences in bone and joint morphometry may be related to age-dependent changes in collagen and extracellular matrix (ECM) organization, although this was not rigorously investigated in the current study.

Several ligands, including *Fgf2, Fgf9*, and *Fgf18*, are expressed in the perichondrium, and both FGF9 and FGF18 regulate chondrocyte hypertrophy in the growth plate by signaling to FGFR3 in chondrocytes (23). Activating mutations of FGFR3 cause achondroplasia, the most common skeletal dysplasia resulting in shortened and bowed lower limbs (40,41). Children with achondroplasia exhibit generalized joint hypermobility and weak muscle tone (42). It is possible that knockdown of *Fgfr1* and *Fgfr2* in our study led to a compensatory upregulation of *Fgfr3*, as has been demonstrated in previous ablation studies using skeletal promoters (Osx-Cre) (25).

This work is not without its limitations. Here, we studied the effect of tendon and attachment-specific ablation of FGF signaling by knocking out only two of the four FGF receptors, *Fgfr1* and *Fgfr2*. Here, we compared the structural but not molecular outcomes associated with FGFR deletion. Although ScxCre *Fgfr1/Fgfr2* DKO adult mice had shortened bones, we do not expect this to be caused by primary growth plate disruptions, as ScxCre is not expressed in proliferating or hypertrophic chondrocytes of the primary growth plate (8). Instead, such differences in bone length may be caused restricted growth plate expansion due to increased expression of FGF9/FGF18 and activation of FGFR3 (25). Even though DKO mice were smaller on average, the effect of partial FGFR signaling ablation resulting in enlarged bone ridges remains noteworthy.

In summary, we have shown that FGF signaling is an important regulator of bone shape and superstructure growth in the adult skeleton. FGF signaling may also play a role in cell survival and matrix remodeling at the tendon and ligament enthesis. The targeted disruption of FGF signaling in tendon and attachment progenitors led to phenotypic changes in bone, joint, and enthesis morphology, including increased bone apposition and cell death at interfaces, underscoring its role in musculoskeletal growth and homeostasis.

## Acknowledgements

This research was supported by the Eunice Kennedy Shriver National Institute of Child Health & Human Development of the National Institutes of Health under award numbers R01AR079367 (MLK), K12HD073945 (MLK), R03HD094595 (MLK), and R01HD049808 (DMO); an Institutional Development Award (IDeA) from the National Institute of General Medical Sciences of the National Institutes of Health under award number P30GM103333; the National Institute of Arthritis and Musculoskeletal and Skin Diseases under award numbers P30AR057235 (WUSTL Musculoskeletal Research Center) and P30AR069620 (Michigan Integrative Musculoskeletal Health Core Center), and the University of Delaware Research Foundation (MLK). EG was supported by the University Doctoral Fellowship Award, ALS was supported by Delaware Space Grant Consortium, and EP was supported by the University of Michigan UROP. Microscopy access was supported by grants from the NIH-NIGMS (P20 GM103446), the NSF (IIA-1301765), and the State of Delaware. Special thanks to Shannon Modla and Jean Ross for assistance with electron microscopy, and to Christopher Price, PhD, for assistance with joint space measurements from microCT.

